# Limited Impact of Schistosome Infection on *Biomphalaria glabrata* Snail Microbiomes

**DOI:** 10.1101/2024.10.18.619126

**Authors:** Stephanie C. Nordmeyer, Timothy J.C. Anderson, Frédéric D. Chevalier, Winka Le Clec’h

## Abstract

**Background:** The microbiome of disease vectors can be a key determinant of their ability to transmit parasites. Conversely, parasite infection may modify vector microbiomes. We are exploring the interactions between the *Biomphalaria glabrata* snail microbiome and the blood fluke *Schistosoma mansoni*, responsible for an estimated 200,000 human deaths each year. Snail hosts vary in their susceptibility to schistosome parasites, and the underlying mechanisms driving this variation are not fully understood. We have previously shown that the snail hemolymph (i.e., blood) and organs harbor a diverse microbiome. Here we investigate the impact of schistosome infection on snail microbiomes, hypothesizing that invading schistosomes can alter the snail microbiomes in both composition and abundance over the course of infection, as developing schistosome parasites are in close contact with the host tissues.

**Result:** We generated cohorts of uninfected and *S. mansoni* infected snails. We collected snail hemolymph and hepatopancreas (i.e., liver) at 8 timepoints during the pre-patent and patent periods of schistosome infection. We quantified bacterial density using qPCR and profiled the microbiome composition of all samples by sequencing the V4 region of the 16S rRNA. Schistosome infection had surprisingly no effect on bacterial density and limited effect on the microbiome composition, affecting mainly the hemolymph during the pre-patent period (at day 7 and 21). Organ and hemolymph microbiomes were relatively stable overtime for both infected and uninfected snail cohorts. The *sample type* (hemolymph, hepatopancreas) was the major driver of the differences observed in microbiome composition.

**Conclusions:** The limited impact of schistosome infection on the host snail microbiomes might be explained by the long-term interaction of the two partners and the fact that parasite fitness is closely dependent on host fitness. Further investigations into the interactions between snails, their microbiomes, and schistosome parasites are essential for developing strategies to disrupt the parasite lifecycle and, consequently, schistosomiasis transmission.

## Introduction

The microbiome plays numerous roles in host fitness and functions, including in metabolism [1–3], immune system development and homeostasis [4,5], and protection against infections or diseases [4,6]. While host microbiomes have been shown to impact invading pathogens [7,8], pathogen infection can also influence the composition of the host microbiome [9,10]. However, the impact of pathogens on the host microbiome may vary significantly depending on whether the pathogens are transient or engage in long-term interactions with the host (i.e., parasitic relationship), especially when pathogen fitness is closely tied to host fitness. The effect of parasites on the host microbiomes has mainly been investigated in vertebrates, with a strong emphasis on gut microbiomes [11]. While parasites often use invertebrate vectors for transmission, their impact on vector microbiomes has received comparatively less attention. The time parasites spent in their vectors, along with the diversity of vectors (insects, crustaceans, mollusks, etc.) and their associated microbiomes, could provide valuable insights into the dynamics of host microbiomes during long-term interactions. A better understanding of these interactions could open new avenues for controlling parasite transmission, as demonstrated by the use of *Wolbachia* bacteria in mosquitoes to manage Dengue virus [12,13].

*Biomphalaria* snails are an excellent model to understand the impact of parasites on host microbiomes. These snails have diverse microbiomes [14,15] and are vectors of a wide range of parasites [16,17], which colonize their host for weeks [18]. Among these parasites, the schistosome blood flukes are of significant biomedical importance, infecting 230 million people in tropical and subtropical areas and responsible for approximately 200,000 deaths per year [19]. Infected people release parasite eggs from which miracidia larvae hatch [20]. These miracidia then actively search for a snail host and, once found, penetrate its head foot. The parasite then develops during several weeks within its host. This development is divided into a pre-patent and patent period [21]. The pre-patent period corresponds to the first four weeks of infection, where the miracidium transforms into primary sporocysts at the site of penetration, and starts producing second generation sporocysts that will be released after 10-11 days [22]. The secondary sporocysts then migrate through snail organs to the hepatopancreas (liver-like organ) and ovotestis [23]. These sporocysts will reside within these organs and, when mature, will produce either additional sporocysts or cercariae, the free-swimming larvae infecting humans. The cercariae are shed from the snail host by breaking through the snail tissues [23]. The start of cercarial shedding marks the beginning of the patent period, which occurs around the 4^th^-5^th^ week of infection.

During its development within its snail host, parasites are exposed to the snail microbiomes, which are not just diverse but also vary across organs and tissues [14,15]. For instance, the hepatopancreas where parasites grow, and the hemolymph (i.e., blood) which bathes organs and parasites, have dramatically different microbiomes [15]. Most microbiome studies of *Biomphalaria* and other parasite-transmitting snails have focused on whole snails [24–29]. However, our recent characterization of hemolymph, organ and whole snail microbiomes revealed that whole snail microbiomes are composite and do not accurately represent the microbiome compositions of the different part of the snail [14,15]. Whole snail microbiomes could, therefore, mask subtle changes or generate spurious shifts in microbiome diversity and composition.

Only a few studies have investigated the impact of schistosome and other trematode infections on snail microbiome. Portet et al. 2021 characterized the microbiomes of whole snails exposed to schistosomes at three time points during the pre-patent period (day 1, 4 and 25 after exposure) which did not reveal disturbance during this period [24]. McCann et al. 2024 examined the whole snail microbiomes of field collected *Galba truncatula* and found differences between uninfected snails and those infected with liver fluke’s larvae [29]. However, the impacts of parasites on specific tissues and organs during the pre-patent and patent periods is unknown.

To better understand the dynamics within relevant snail organ compartments, we characterized the impacts of *S. mansoni* infection on the microbiome of *B. glabrata* hemolymph (to which parasites are exposed) and hepatopancreas (where parasites reside) over the course of infection. We performed weekly collections of infected and non-infected snails over a six-week period, quantifying the bacterial burden and characterizing the microbiome composition of sampled tissues. Our results showed a limited and specific impact of schistosome infection on the composition of the hemolymph microbiome.

## Materials and Methods

### Snail infection

We used *Biomphalaria glabrata* snails from the inbreed line 26 [Bg26, derived from 13-16-R1 line [30]] to limit the impact of the host genetic variation on our study. Snails were reared in 10-gallon aquariums with well water at 26-28°C on a 12-hour light cycle and fed *ad libitum* on green lettuce [21]. All snails used in the study had a shell diameter between 5-7 mm and were randomly sampled from two aquariums and acclimated in trays (55 snails per tray) 7 days before exposure to parasites. In addition to being inbred, Bg26 snails are highly susceptible to *Schistosoma mansoni* miracidia from SmLE population [31]. We prepared miracidia by emulsifying livers of two Syrian golden hamsters as previously described [31]. We exposed Bg26 snails (n = 110) to 5 SmLE miracidia in 1 mL freshwater in a 24-well plate under artificial light on D0 overnight [31]. We prepared a separate cohort of 110 control snails into plates to undergo the same procedure of exposure without miracidia. The next morning, we moved snails to trays with well water and green lettuce. Each cohort had 2 trays, and each tray had 55 snails (Control trays: CA, CB; Exposed trays: EA, EB). We kept trays in a temperature-controlled room at 26-28°C on a 12-hour light cycle. During normal maintenance of laboratory infected snails, we cover the trays with a black plexiglass lid starting on week 3 post-exposure to parasite to prevent cercarial shedding from infected snails. To better mimic natural field conditions, the experimental snails were covered only with a clear plexiglass lid for the duration of the experiment. Well water and lettuce were replaced daily. Snails were visually assessed for infection during dissection, starting at day 21 by looking at the presence of schistosome sporocysts or cercariae in ovotestis and hepatopancreas. The infection rate for D7 and D14 was estimated based on the overall observed infection rate from D21 to D39.

### Hemolymph and hepatopancreas collection

Snails were collected at 8 time points during the course of infection, on D0 (prior to exposure), D7, D14, D21, D28, D35, D37 and D39. For each collection, 3-4 snails were sampled from each tray (6-8 snails per cohort; 12-16 snails per timepoint). We sampled 500 µL of water from each tray at these timepoints (environmental samples). Each snail was then processed individually. We wiped the shell 3 times with 70% ethanol for disinfection. Hemolymph was collected by heart puncture using a sterile 1 mL syringe and 23.5-gauge needle [14]. We then gently crushed the snail shell between 2 sterile microscope slides and removed shell pieces with sterilized tweezers. We collected the hepatopancreas (liver) and rinsed it three times with 1 mL of sterile water. Each collected sample was immediately placed in a sterile 1.5 mL pre-cooled microtube stored on dry ice.

### 16S rRNA gene library preparation and sequencing

We randomized all samples prior to both DNA extraction and triplicate PCR assays for library preparation to minimize any potential batch effects. Samples were randomized based on factors including tray, snail number, sample type (water, hemolymph, hepatopancreas) and day of collection. We extracted DNA from the snail samples and tray water samples using the DNeasy Blood and Tissue Kit (Qiagen) following the manufacturer’s protocol with minor adjustments [14]. “Kitome” controls were included to check for any potential contamination in the DNA extraction kit [32]. We used 40 µL of hemolymph and environmental water sample for each DNA extraction. We manually homogenized the hepatopancreas organs using sterile micropestles prior to extraction. Samples were incubated for one hour at 56 °C in a water bath. We recovered the final gDNA in 50 µL of elution buffer. The 16S rRNA gene libraries were prepared in a biosafety cabinet using sterile equipment and materials to avoid contamination. We performed triplicate PCR for each sample to limit potential amplification biases introduced during the PCR reaction. The 16S rDNA V4 region was amplified using primers (515f and 806rB) from the Earth Microbiome Project [33]. Each 515f primer was barcoded for sample identification. We followed the methods described in Chevalier et al. 2020 [14]for generating the 16S V4 libraries, with some modifications: For each PCR reaction mix, we used 1 µL sterile water, 5 µL AccuStart II PCR Supermix, 1 µL of each primer at 2 µM, and 2 µL of DNA template. Libraries were purified using KAPA Pure Beads and quantified using Picogreen assay (Invitrogen) following manufacturer’s protocol. Two equimass library pools (124 samples each) were made using 30 ng of each library. These pooled libraries were quantified by qPCR with the KAPA library quantification kit, following manufacturer’s protocol, and sequenced by Admera Health on an Illumina MiSeq platform (250-bp paired-end sequencing). Raw sequencing data are accessible from the NCBI Sequence Read Archive under BioProject PRJNA1171869.

### Measure of bacterial density

We measured bacterial density within the hemolymph by amplifying the V4 regions of the 16S rRNA gene using primers 515f and 806rB (without adaptors and barcodes) by qPCR as previously described [14]. We determined absolute 16S rRNA gene quantities by using standard curves of six 10-fold dilutions (6×10^1^ to 6×10^7^) of purified 16S rRNA gene PCR product from *E. coli*. Results were normalized by the amount of hemolymph used in the DNA extraction (40 µL) and the qPCR dilution factor (1/10^th^ dilution). Samples with a normalized quantity above 500,000 copies (outliers in the distribution of the data) were excluded from the analysis.

### Bioinformatic and statistical analysis

We performed sequence processing and analysis using QIIME2 (v2021.4) [34] and R (v4.3.1) [35]. Scripts used for sequence processing and analysis are available in a Jupyter notebook (https://github.com/snordmeyer/S.mansoni-infected-B.glabrata-microbiomes). We processed the sequencing data as previously described [14]. Briefly, we denoised and clustered the sequences into ASVs using the dada2 module with a max expected error of 5. We denoised only the forward reads due to the low quality of the reverse reads. We determined the taxonomy of the ASVs using the SILVA database (release 132). We blasted ASVs with unassigned taxonomy against the NCBI nt database using megablast from blast+ to identify eukaryotic contaminants for removal in downstream analysis. We aligned the ASVs and masked the highly variable positions using the mafft and mask commands from the alignment module to build a phylogenetic tree. Finally, we built and rooted a tree from the alignment using the fasttree and midpoint-root commands.

We imported the QIIME2 files in R using the *qiime2R* package (v0.99.6) and converted them into *phyloseq* objects using the *phyloseq* package (v1.46.0). We assessed alpha diversity using the number of observed ASVs and Simpson Evenness using the package *microbiome* (v1.24.0). None of the alpha-diversity data followed a normal distribution (Shapiro-Wilk test, p-value < 0.05). We therefore used a non-parametric (Wilcoxon or Kruskal-Wallis and Dunn’s post-hoc) test for the statistical comparison of the data. We assessed beta diversity on rarefied data using R packages *vegan* (v2.6-4) and *pairwiseAdonis* (v0.4.1) to perform beta-dispersion and PERMANOVA tests including factors *cohort* and *day* with 1,000 permutations. We performed rarefaction and statistical tests for each sample type (hemolymph, hepatopancreas, water). We assessed the data for homogeneity of variances by computing the centroid distances and assessing differences using the TukeyHSD post-hoc test. Additive and interaction linear mixed models including fixed factors (*cohort, sample type, day*) and random factors (*snail_ID, tray*) were assessed for each alpha- and beta-diversity index using packages *stats* (v4.3.1), *lme4* (v1.1-34) and *lsmeans* (v2.30-0). We used the Bayesian Information Criterion (BIC) scores for each statistical model tested to determine the best fitting models to report.

We also assessed bacterial density data using the non-parametric Wilcoxon or Kruskal-Wallis and Dunn’s post hoc tests for statistical comparisons.

## Results

### Experimental design and library statistics

We sampled the hemolymph and hepatopancreas of *B. glabrata* snails uninfected and infected with *S. mansoni* parasites over the course of 39 days (Figure 1). For each of the 8 timepoints, we collected 6-8 uninfected (control, unexposed cohort) and infected (exposed cohort) snails. The snail infection rate was 100% after visual assessment of the hepatopancreas during dissection starting D21 (presence of daughter sporocysts) and onwards (presence of cercariae). We assumed snails from prior D21 were also 100% infected in the *S. mansoni* exposed cohort. Such high infection rate was expected given the high susceptibility of Bg26 snails to our SmLE population. Survival rates of snails not sampled for microbiome analysis were similar between replicate control or exposed *S. mansoni* treatments [(CA: 11/22, 50% survival; CB: 15/22, 68.18% survival; Log Rank test = 2.2, p-value = 0.1) (EA: 3/24, 12.5% survival: EB: 2/24, 8.33% survival; Log Rank test = 0, p-value = 1)] (SuppFigure1). However, *S. mansoni* infection had a significant impact on survival, with control snails having a higher survival rate than the exposed snails, as expected [36] (Log Rank test = 13.8, p-value = 2e-04). We observed marked increase in mortality in the exposed snails around D30.

**Fig. 1.**
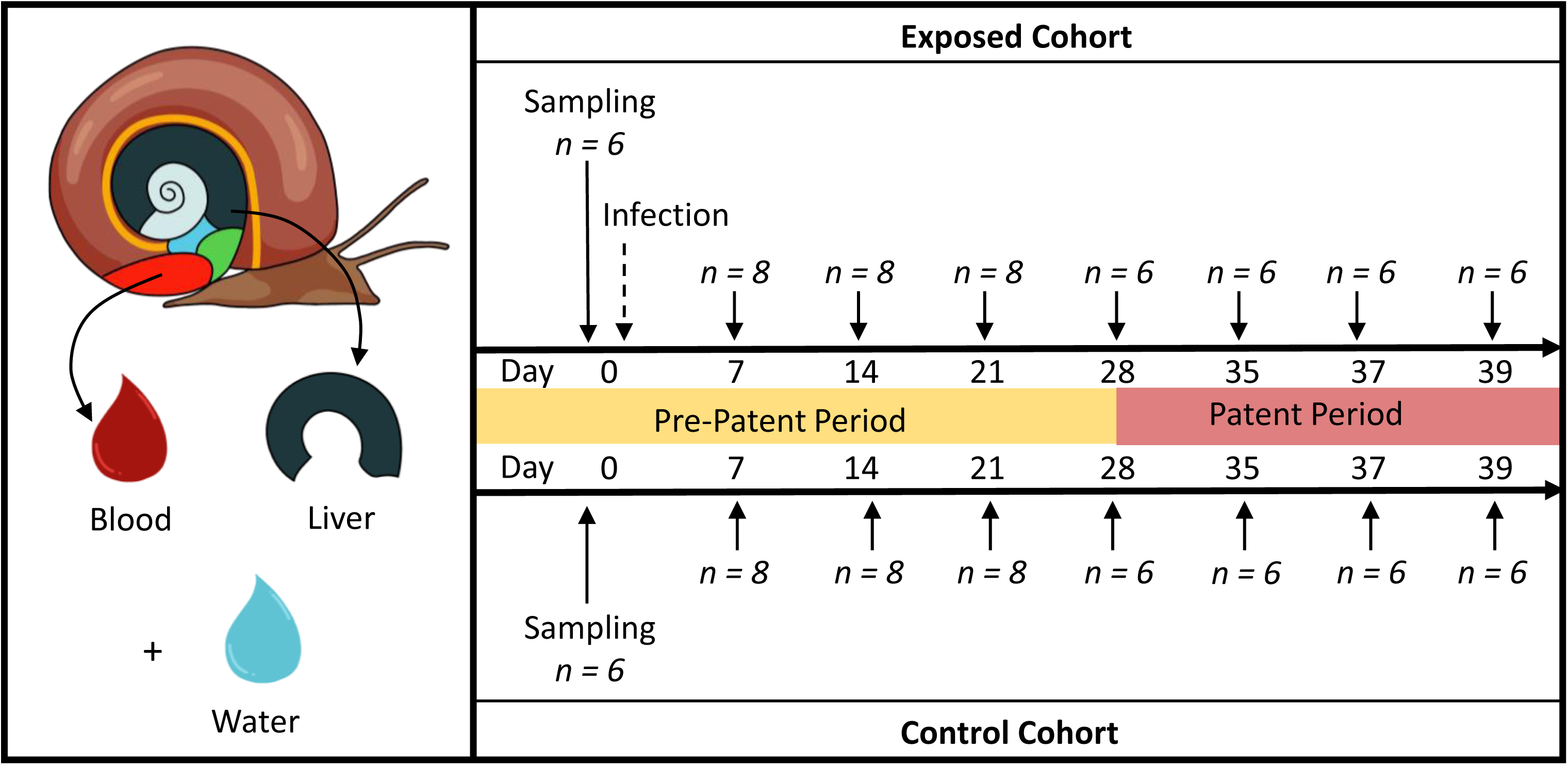
Experimental Design. *Biomphalaria glabrata* Bg26 snails were randomly assigned to the exposed (n = 55) or control (n = 55) cohort. Six snails were sampled on D0 prior to miracidial or mock exposure in both the exposed and the control cohorts. Snails were then sampled weekly, through the pre-patent (n = 8 snails per cohort – D7 to D21) and patent (n = 6 snails per cohort – D28 to D39) periods. At these timepoints, the hemolymph and hepatopancreas of each snail was collected under sterile conditions, along with an environmental water sample from each tray.

A total of 216 snail samples (108 hemolymph and 108 hepatopancreas) and 32 environmental samples (tray water) were collected and processed for microbiome library preparation. We used high-throughput sequencing of the V4 regions of the 16S rRNA gene with the Illumina MiSeq platform, and obtained a total of 14,428,659 reads from 248 samples with an average of 58,180 ± 1,406 (mean ± S.E.) input reads per sample. After filtering, we had 9,836,712 reads with an average of 39,664 ± 918 filtered reads per sample (SuppFigure2, SuppTable1). We identified 4,923 ASVs and removed ASVs matching mitochondria and chloroplasts (82 ASVs, 1.7% of ASVs) or eukaryotes (149 ASVs, 3.1% of ASVs). Among the 4,692 ASVs retained, 43.27% (2,030) had unassigned taxonomy. Rarefaction curves showed that all samples reached a plateau, confirming that our sequencing effort was sufficient to capture all the bacterial diversity present in our samples (SuppFigure3).

### Bacterial density is stable in uninfected and infected snail hemolymph over time

We measured the impact of *S. mansoni* infection on the bacterial density of snail hemolymph by comparing bacterial loads between infected and uninfected snails at each timepoint using qPCR. Infected snails showed stable bacterial density in the hemolymph over time (Kruskal-Wallis test: X^2^ = 13.19, df = 7, p = 0.0676; Figure 2). Bacterial density measured in the hemolymph of the uninfected snail cohort also remained relatively stable, with the only significant difference observed between days 28 and 37 (Kruskal-Wallis test: X^2^ = 17.171, df = 7, p = 0.0163, comparison between D28 vs. D37: p = 0.0482; Figure 2). In addition, there was no significant differences at any given timepoints between the infected and uninfected cohorts (Table 1; Wilcoxon test: W= 10 to 27, p = 0.27 to 1). Overall, schistosome infection did not impact the bacterial density in the hemolymph. In addition, hemolymph microbiome showed a strong temporal stability suggesting a tight control of the microbiome by the snail.

**Fig. 2.**
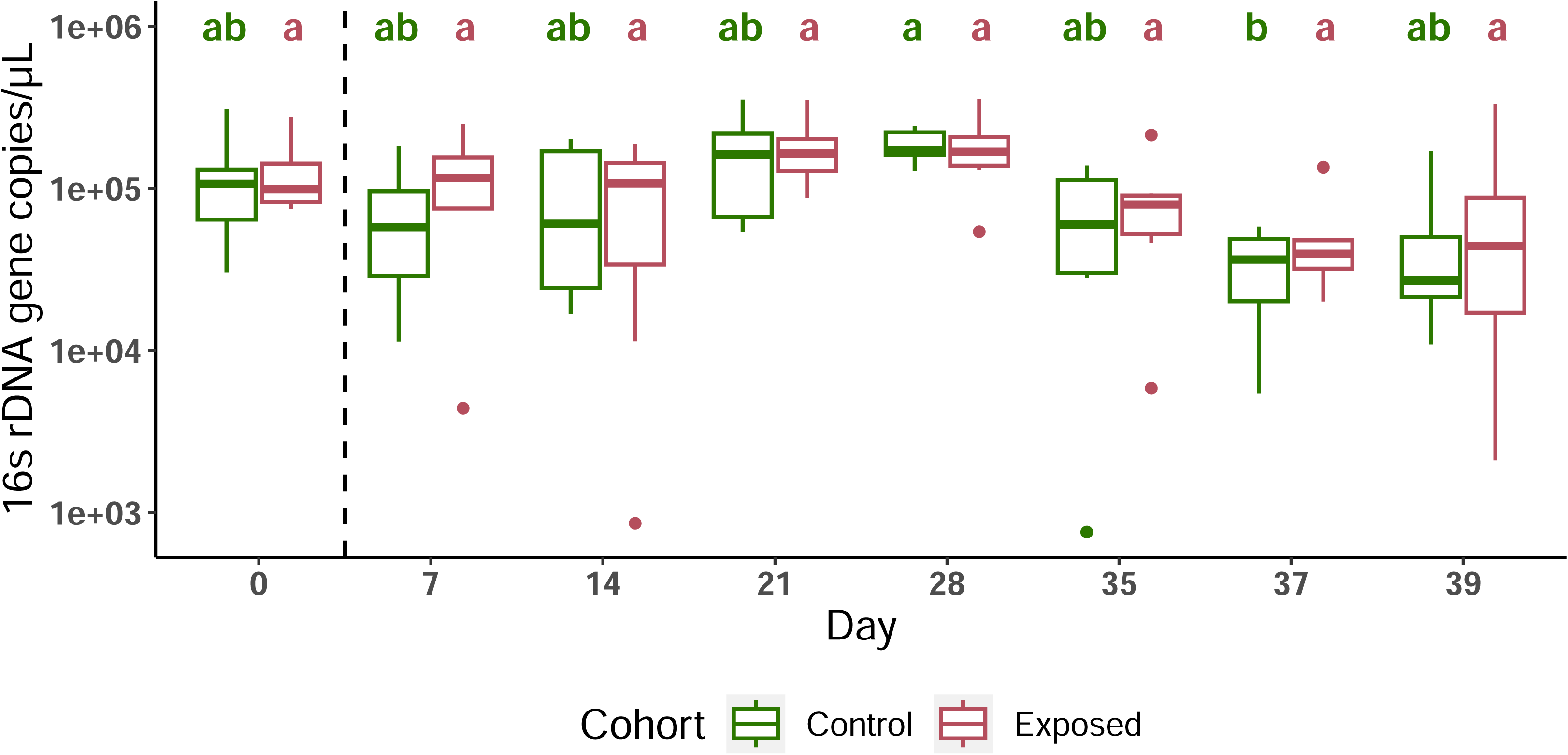
Hemolymph bacterial densities between control and schistosome-exposed snails. The longitudinal evolution of the number of 16S rRNA gene copies per microliter of snail hemolymph (D0 to D39, measured by qPCR), showed no striking difference between cohorts at any of the timepoints. The dashed line marks period before snails were exposed to miracidia (D0), and after snails have been exposed (D7 onwards). Statistical differences within cohorts at each of the timepoints were assessed using Kruskal-Wallis and Dunn’s post-hoc tests for non-parametric data. Letters generated from Dunn’s post-hoc tests are shown above the boxplots. Groups under the same letter are not statistically different. Control: green, Exposed: red.

**Table 1.** Bacterial Density statistics. Wilcoxon testing of significant differences in bacterial density between infected and uninfected snails at each timepoint. *W = Wilcoxon test statistic*.

| Day | W | p-value |
| --- | --- | --- |
| 0 | 12 | 1.000 |
| 7 | 21 | 0.270 |
| 14 | 27 | 0.798 |
| 21 | 21 | 1.000 |
| 28 | 16 | 0.927 |
| 35 | 17 | 0.936 |
| 37 | 14 | 0.575 |
| 39 | 10 | 0.749 |

Similarly, we attempted to assess the impact of schistosome infection on the bacterial density of the hepatopancreas organ, where schistosome parasites reside. However, we were unable to amplify the snail reference gene (*piwi4*) in the majority of the samples, preventing normalization of the data.

### Schistosome parasite infection has minimal impact on the microbiome diversity of its snail host

#### 1. α-Diversity

We assessed α-diversity by measuring the total number of observed ASVs (species richness), Simpson evenness (species evenness) and Faith’s phylogenetic diversity (phylogenetic richness) of each snail hemolymph and hepatopancreas sample.

##### Species richness

We first assessed how much variation in species richness was explained by each factor using a fixed effect model (*observed ASVs ∼ cohort + day + sample type*). *Sample type* (snail tissues [hemolymph, hepatopancreas], water) explained the majority of the variation in microbiome richness, followed by *day* (Table 2). Next, we explored which factors specifically shaped species richness in each snail tissue sample type using a second fixed effect model (*observed ASVs ∼ cohort + day*). *Day* was the only significant factor for both hemolymph and hepatopancreas while the *cohort* factor (exposed to *S. mansoni* parasites or control) had no significant effect (Table 2).

**Table 2.** Alpha-Diversity linear mixed model results. ANOVA results (F statistics and associated P value) on the additive models used on the alpha-diversity metrics: observed ASVs, Simpson evenness and Faith’s phylogenetic diversity. The linear mixed models included all three sample types (hemolymph, hepatopancreas, environmental water) or were run individually for hemolymph and hepatopancreas samples.

|  | Observed ASVs |  | Simpson Evenness |  | Phylogenetic Diversity |  |
| --- | --- | --- | --- | --- | --- | --- |
| *All tissues* | F value | P value | F value | P value | F value | P value |
| **Cohort** | 1.4155 | 0.2353 | 1.0097 | 0.316 | 2.0124 | 0.1573 |
| **Sample Type** | 78.4187 | <2.2e-16 | 10.5302 | 0.000042 | 110.7939 | <2.2e-16 |
| **Day** | 8.3457 | 4.136E-09 | 0.736 | 0.6416 | 7.0648 | 1.154E-07 |

| *Hemolymph* |  |  |  |  |  |  |
| --- | --- | --- | --- | --- | --- | --- |
| **Cohort** | 2.8044 | 0.097163 | 2.2443 | 0.1373 | 3.1122 | 0.080792 |
| **Day** | 3.6561 | 0.001498 | 1.0121 | 0.4275 | 2.8392 | 0.009688 |

| *Hepatopancreas* |  |  |  |  |  |  |
| --- | --- | --- | --- | --- | --- | --- |
| **Cohort** | 1.9106 | 0.17 | 3.0444 | 0.08412 | 1.6175 | 0.20643 |
| **Day** | 2.6328 | 0.01545 | 1.3688 | 0.22697 | 2.3729 | 0.02765 |

We only observed a single timepoint with a significant difference in the number of ASVs between the hemolymph of control and exposed snails, on D21 (Wilcoxon test: W = 55, p = 0.018) with control snails having a higher number of ASVs (Figure 3A). Water showed extensive variation in observed ASVs over time (range: 92 – 489, mean: 233.406), however this variation did not appear to affect the number of ASVs in the snail hemolymph or hepatopancreas. The species richness index showed a significantly higher number of ASVs in the hemolymph compared to the hepatopancreas and environmental water (Figure 3A). We found approximately 2 times more ASVs in the hemolymph (262.407 ± 8.422) compared to the hepatopancreas (142.564 ± 5.341) samples. The average number of observed ASVs for water samples (233.406 ± 19.283) was comparable to that of hemolymph; however, the number of observed ASVs in water showed dramatic variations over time compared to the hemolymph samples (SuppTable2).

**Fig. 3.**
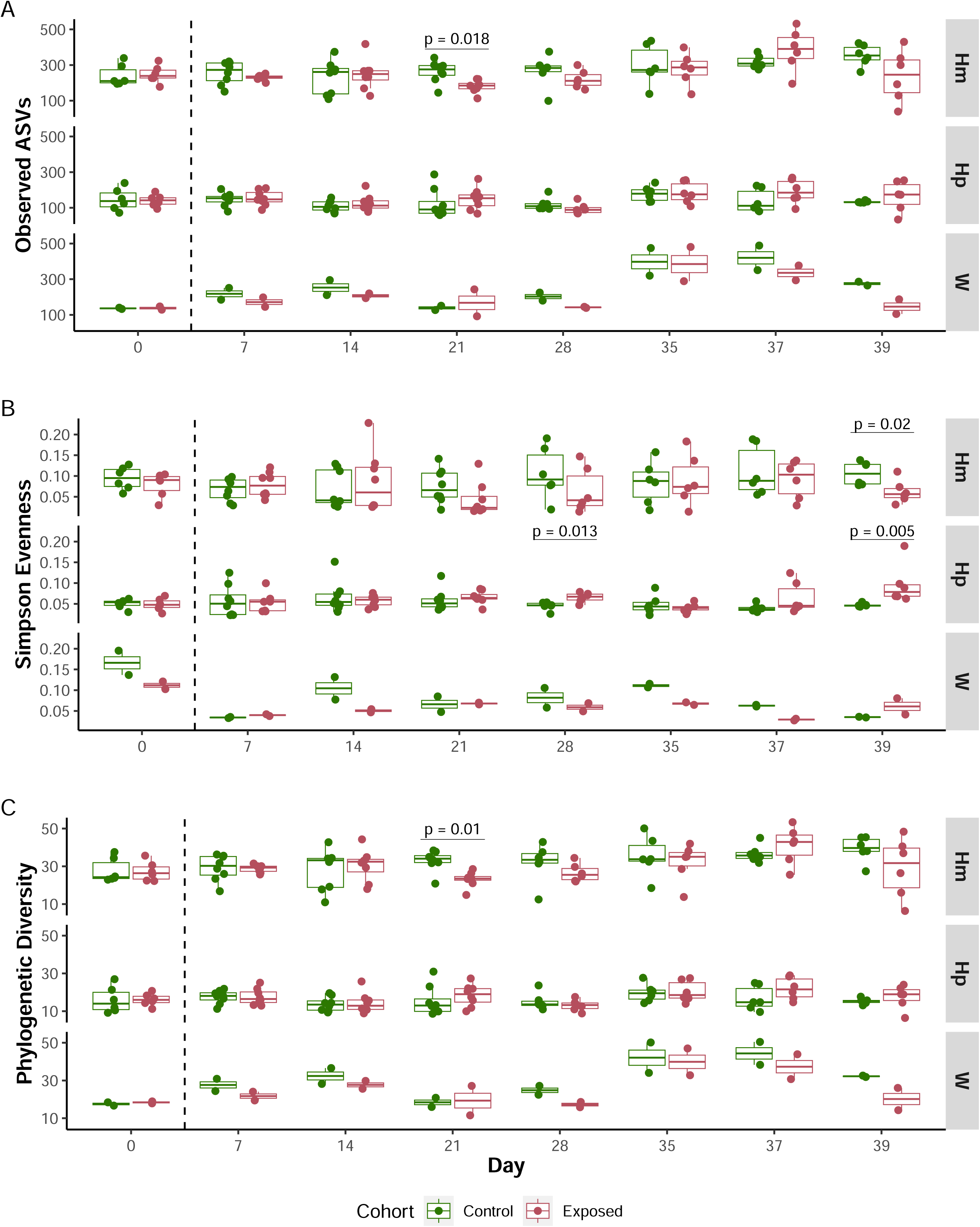
Longitudinal comparison of alpha-diversity metrics between control and schistosome-exposed snails. Boxplots showing values for (A) observed ASVs, (B) Simpson Evenness, and (C) Faith’s Phylogenetic Diversity between control (green) and exposed (red) snail cohorts at each of the 8 timepoints, and for each sample type (Hm: hemolymph, Hp: hepatopancreas, W: water). The longitudinal analysis of the observed ASVs and the Faith’s PD metrics reveals specific timepoints where schistosome infection significantly impacts the microbial composition of hemolymph (D21 and D39) or hepatopancreas (D28 and D39). The dashed line marks period before snails were exposed to miracidia (D0), and after snails have been exposed (D7 onwards).

##### Species evenness

We measured the distribution of ASVs using the Simpson Evenness index (Figure 3B). The s*ample type* factor explained all the observed variation in species evenness, based on the best fitting model for this index (*evenness ∼ cohort + day + sample type*) (Table 2).

This community evenness was low across all samples (Figure 3B), exhibiting a wider range in the hemolymph (range: 0.014 - 0.228) compared to the hepatopancreas (range: 0.023 - 0.19) (Fligner-Killeen test: X^2^ = 12.914, df = 1, p = 0.0003), as previously observed [15]. In the hemolymph, we observed a significant decrease in species evenness only in exposed snails compared to control on day 39 (Wilcoxon test: W = 33, p.adj = 0.02) (SuppTable3). In the hepatopancreas, significant differences in evenness between cohorts were observed on day 28 and 39 (D28: Wilcoxon test: W = 2, p.adj = 0.013; D39: Wilcoxon test: W = 0, p.adj = 0.005).

##### Phylogenetic diversity

We estimated the phylogenetic differences between microbial communities using the Faith’s phylogenetic diversity index. Using the same fixed model as for species richness and evenness, we found that the *sample type* factor, again, explained most of the variation observed in the phylogenetic diversity (Table 2). When we applied the model to each snail tissue separately, the *day* factor significantly explained the variation observed in both the hemolymph and hepatopancreas samples.

Similar to species richness, only one timepoint – day 21 in the hemolymph – showed significant differences between cohorts (Wilcoxon test: W = 57, p.adj = 0.01) (SuppTable3). At day 21, control snails had significantly higher microbial phylogenetic diversity (32.381 ± 1.154) in their hemolymph compared to exposed snails (29.562 ± 1.238) (Figure 3C). The phylogenetic diversity between tissues (Figure 3C) mirrored the observed ASVs diversity, with water samples showing similar diversity levels to hemolymph (SuppTable3), and hepatopancreas samples exhibiting the lowest diversity.

#### 2. *β*-Diversity

We conducted the β-diversity analysis on hemolymph and hepatopancreas separately due to the strong effect of the *sample type* factor observed in the α-diversity analysis. To ensure that differences observed by PERMANOVA were due to group distance rather than group dispersal, we tested for the homogeneity of variances using a β dispersion analysis. Nearly all of the hemolymph and hepatopancreas samples showed similar group dispersion between cohorts, with the exception of three hemolymph comparisons and a maximum of four hepatopancreas comparisons (SuppFigure4).

We then explored differences between sample groups using Bray-Curtis and weighted UniFrac distances. Both metrics quantify dissimilarity between samples, with Bray-Curtis accounting for species abundance, and weighted UniFrac also considering the phylogenetic relatedness of ASVs. We tested the effect of the factors *cohort* and *day* on the hemolymph groups using PERMANOVA analysis (*distance ∼ cohort + day*) (Table 3). Both *cohort* and *day* significantly explained the variation observed with the Bray-Curtis distance (Cohort: PERMANOVA: F = 3.9705, p = 0.0009; Day: PERMANOVA: F = 4.6403, p = 0.0009) (Figure 4A, SuppFigure5A). In contrast, neither factor *cohort* nor *day* explained the overall variation observed with weighted UniFrac distance (Cohort: PERMANOVA: F = 0.5674, p = 0.5215; Day: PERMANOVA: F = 1.6243, p = 0.0959) (Figure 4B, SuppFigure5B). These results suggest that parasite infection may alter the relative abundance of certain taxa without significantly affecting the broader evolutionary composition of the microbiome. Additionally, we found that the hemolymph microbiome composition was significantly different between infected and uninfected snails, and within the infected cohort at days 7, 21, 37 and 39 with Bray-Curtis, and days 7 and 21 with weighted UniFrac. This suggests specific microbiome changes occurring at defined developmental stages of the schistosome parasite within its snail host.

**Fig. 4.**
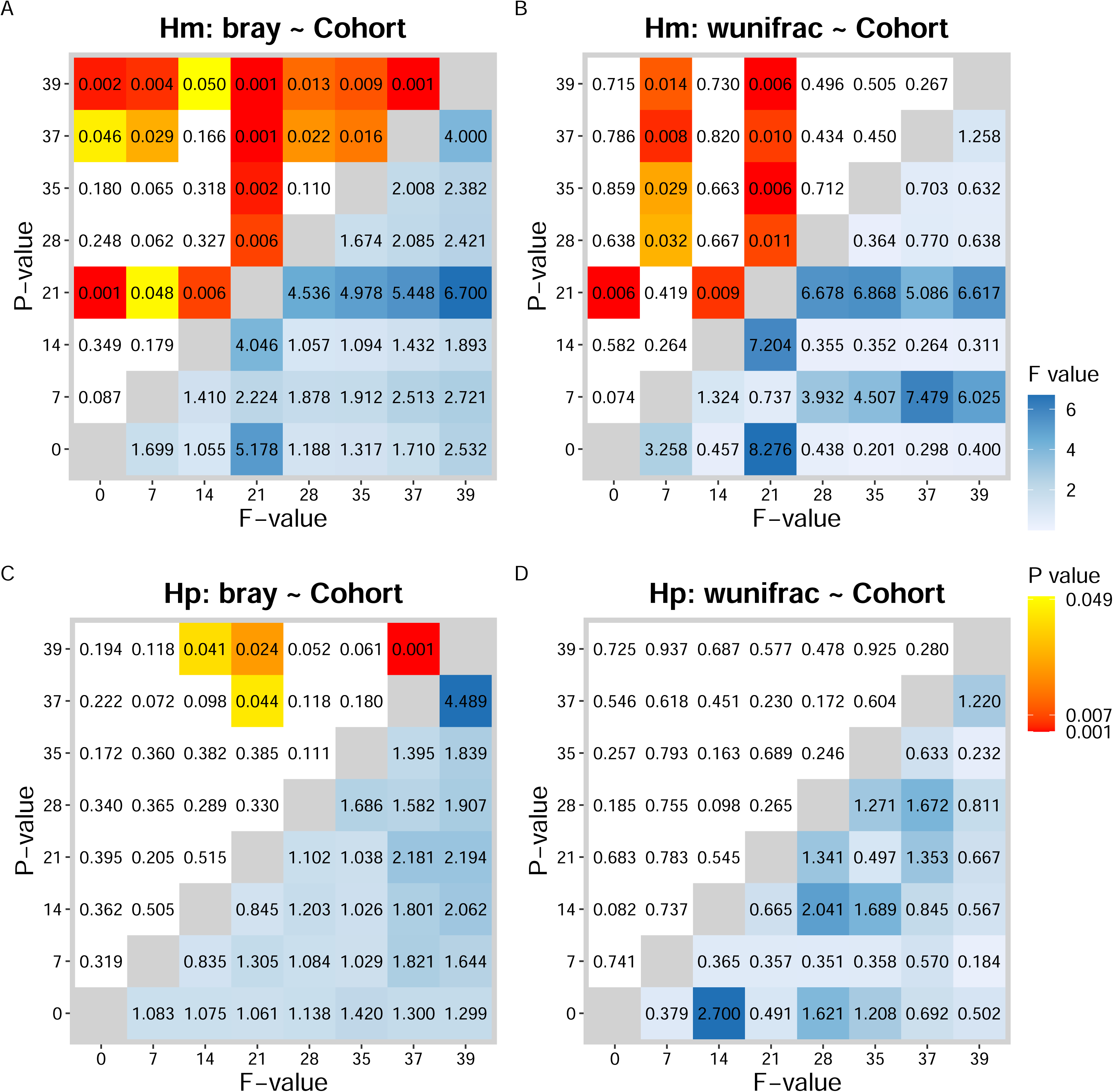
Longitudinal comparison of beta-diversity metrics between control and schistosome-exposed snails, for hemolymph and hepatopancreas. Heatmaps depicting the statistical significance between microbiome sampled on different days determined by using a pairwise post-hoc test on the PERMANOVA models for (A-B) hemolymph (Hm) and (C-D) hepatopancreas (Hp), using the (A, C) Bray-Curtis and (B, D) weighted UniFrac metrics. The lower triangle shows pairwise F-values, the upper triangle shows p-values, and the days compared are marked on the axes. The hemolymph microbiome varied significantly between infected and uninfected snails, especially at D7 and D21 for both metrics, showing stage-specific changes during schistosome development. The hepatopancreas microbiome showed minimal impact from infection.

**Table 3.** Beta-Diversity PERMANOVA model results. PERMANOVA (Permutational Multivariate Analysis of Variance using Distance Matrices) results on the additive models used on the beta-diversity metrics Bray-Curtis and Weighted UniFrac. Models were run separately for each snail tissue (hemolymph and hepatopancreas).

|  | **Bray Distance ~ Cohort + Day** |  |  |  |  |
| --- | --- | --- | --- | --- | --- |
| *Hemolymph* | **df** | **Sum of Squares** | **R<sup>2</sup>** | **F** | **P value** |
| **Cohort** | 1 | 0.8785 | 0.0293 | 3.9705 | 0.000999 |
| **Day** | 7 | 7.1869 | 0.2398 | 4.6403 | 0.000999 |
| **Residual** | 99 | 21.9045 | 0.7309 |  |  |

| *Hepatopancreas* |  |  |  |  |  |
| --- | --- | --- | --- | --- | --- |
| **Cohort** | 1 | 0.3129 | 0.0152 | 1.9500 | 0.038961 |
| **Day** | 7 | 4.4182 | 0.2143 | 3.9336 | 0.000999 |
| **Residual** | 99 | 15.885 | 0.7705 |  |  |

|  | **Weighted UniFrac Distance ~ Cohort + Day** |  |  |  |  |
| --- | --- | --- | --- | --- | --- |
| *Hemolymph* | **df** | **Sum of Squares** | **R<sup>2</sup>** | **F** | **P value** |
| **Cohort** | 1 | 0.00428 | 0.0051 | 0.5674 | 0.5215 |
| **Day** | 7 | 0.08584 | 0.1025 | 1.6243 | 0.0959 |
| **Residual** | 99 | 0.74739 | 0.8924 |  |  |

| *Hepatopancreas* |  |  |  |  |  |
| --- | --- | --- | --- | --- | --- |
| **Cohort** | 1 | 0.00364 | 0.0114 | 1.3002 | 0.25874 |
| **Day** | 7 | 0.03795 | 0.1191 | 1.9374 | 0.01099 |
| **Residual** | 99 | 0.27701 | 0.8695 |  |  |

We performed the same analysis with the hepatopancreas samples. Similarly, both *cohort* and *day* factors explained the variation observed with the Bray-Curtis distance (Cohort: PERMANOVA: F = 1.95, p = 0.0389; Day: PERMANOVA: F = 3.9336, p = 0.0009) (Figure 4C, SuppFigure5C). However, only the *day* factor explained the observed dissimilarities with weighted UniFrac distance (Cohort: PERMANOVA: F = 1.3002, p = 0.2587; Day: PERMANOVA: F = 1.9374, p = 0.01099) (Figure 4D, SuppFigure5D). While we observed some significant differences between days in the hepatopancreas microbiome, there were no consistent patterns at these timepoints. Additionally, we found no significant effect of the *cohort* in the microbiome composition in this tissue. These results suggest that schistosome infection does not significantly impact the microbiome composition of the hepatopancreas.

### Taxonomy

We investigated the impact of *S. mansoni* infection on the microbiome community structure of *B. glabrata* snail hemolymph and hepatopancreas over time in both uninfected and infected snails. The microbiome in snail samples (hemolymph and hepatopancreas) and environmental (water) samples were dominated by Proteobacteria, Bacteroidetes, Tenericutes, and unassigned phyla (Figure 5). While the microbiome composition of snail hemolymph and hepatopancreas was consistent with previous findings [15], the taxonomic composition for the water samples was unexpected. In our previous studies [14,15], water samples were dominated by Actinobacteria, Bacteroidetes and Proteobacteria, whereas Actinobacteria was almost absent in the current study. This difference may be attributed to the water source and/or the frequency of the water change. In the present study, snails were kept in well water changed daily, whereas snails from previous studies were sampled from tanks containing several-months-old well water.

**Fig. 5.**
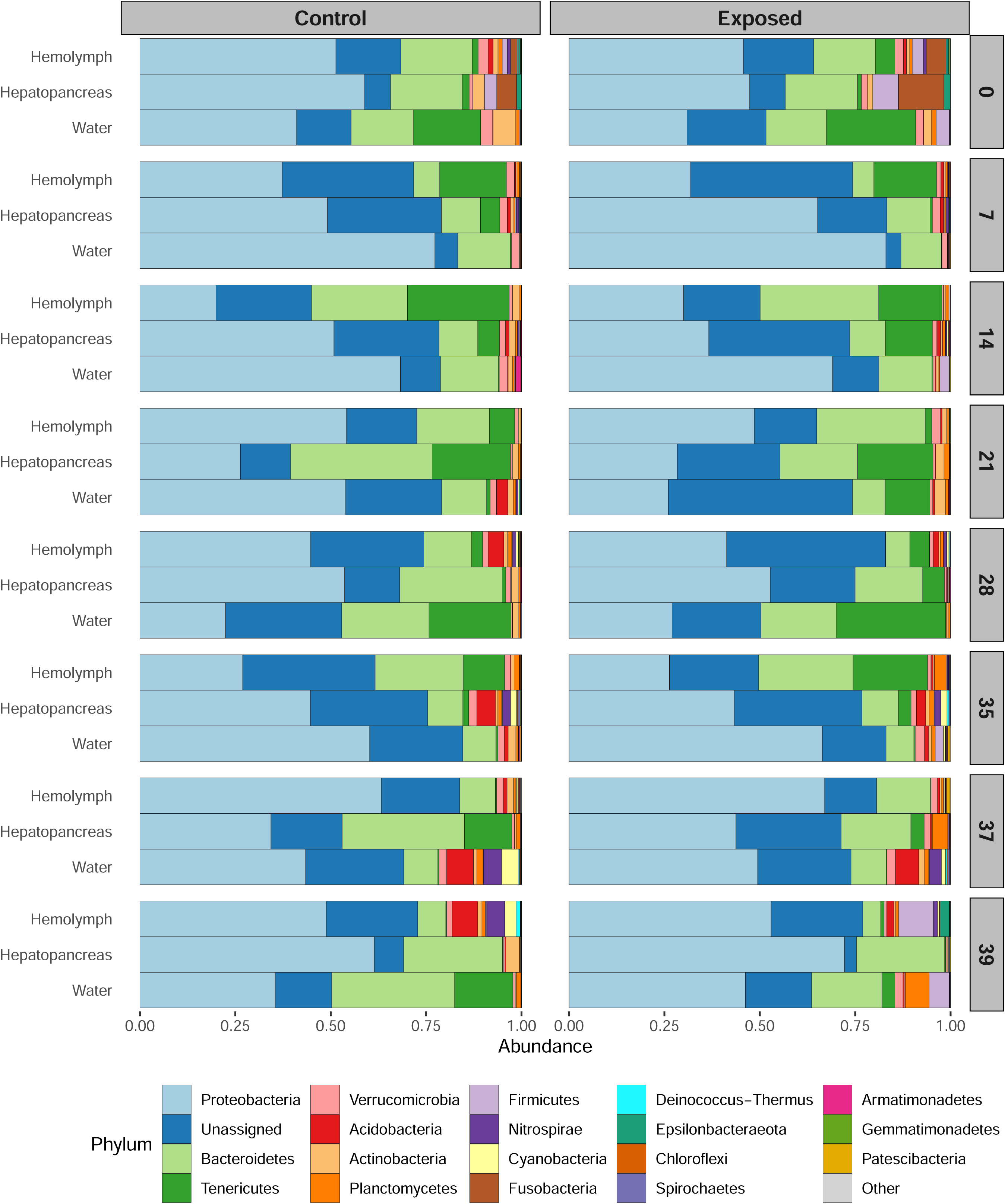
Longitudinal taxonomic diversity between control and schistosome-exposed snails. The relative abundance of the top 20 most dominant phyla is shown for both control (left) and exposed (right) cohorts at each of the 8 timepoints, including snail tissues (hemolymph and hepatopancreas) and environmental water samples. Samples were merged by sample type and day: each bar represents 2 water samples, and 6-8 snail samples for hepatopancreas or hemolymph. The taxonomic composition of both Control and Exposed snails are relatively stable over time, dominated by phyla Proteobacteria, Bacteroidetes, and Tenericutes. We observed significant differences in the microbiome composition between snail tissues (hepatopancreas and hemolymph) and the water environment.

As observed with Bray-Curtis distances, significant differences in microbiome compositions were evident between each sample type. While microbiome compositions remained relatively consistent over time, the proportion of the different phyla exhibited variations. For instance, the proportion of Fusobacteria dramatically decrease in both hemolymph and water after the start of the experiment, for both controls and exposed cohorts. We observed a similar decrease for Actinobacteria in the hepatopancreas. In contrast, Acidobacteria proportions increased in hemolymph, particularly in the control cohorts starting at day 21. A similar increase was observed for Nitrospirae and Cyanobacteria at day 35. Infected snails also showed a dramatic increase in Firmicute on day 39, both in hemolymph and hepatopancreas.

## Discussion

### 1. Limited and specific effect of schistosome on snail hemolymph microbiome during infection

We investigated the impact of schistosome parasite infection on the microbiomes of both the hemolymph and hepatopancreas in *Biomphalaria* snails (Figure 1). We selected these two sample types for three reasons: (i) both harbor diverse and specific microbiomes [15], (ii) the hemolymph bathes the snail’s organs and parasites, and has the greatest microbial diversity [14,15], and (iii) the hepatopancreas is the primary organ where schistosome parasite sporocysts reside and develop [37]. The comparison of microbiomes between control and exposed snails during the pre-patent (before cercariae larvae are produced and shed) and patent periods (when cercariae are produced and released) revealed no significant overall impact of parasitism. The primary difference observed was between *sample type* (hemolymph vs. hepatopancreas), consistent with our previous observations in uninfected snails [15]. This difference in microbial composition of the snail tissues persisted over time, regardless of infection status. We also observed an effect of time (*day* factor), but to a lesser extent.

While infection status had no overall impact on the host microbiome, we observed differences at very specific timepoints. During the pre-patent period, the hemolymph microbiome of infected snails showed differences in composition at day 7 and 21, with the latter likely explained by a lower microbial diversity in infected snails. Day 21 is a critical timepoint in schistosome development: by this stage, schistosome sporocysts have migrated through host tissues and are actively colonizing and multiplying within the snail’s hepatopancreas [37]. During the patent period, we observed differences in community evenness at day 28 for hepatopancreas and at day 39 for both the hemolymph and hepatopancreas. Heavy colonization of the snail hepatopancreas and ovotestis [37,38] and tissue damage due to cercarial shedding could explain these differences. Previous studies using schistosome parasites have shown that the microbiomes of the mammalian [39,40] and the invertebrate hosts [24] can be marginally impacted by infection. Portet *et al.* showed no change in α-diversity of the whole snail microbiome after infections of *B. glabrata* snails (BgBRE) with *S. mansoni* (SmBRE and SmVEN) on days 1, 4 and 25 after primary infection [24]. Our results, focusing on organ and hemolymph microbiomes, indicate similar findings at the beginning of the pre-patent period but suggest specific changes toward the end of this period. This difference could be attributed to the use of SmLE parasites, which are known to have a stronger impact on the snail host [31], or it may have been revealed by the analysis of tissue-specific microbiomes that were otherwise masked by the use of whole snails [24].

We observed no significant differences in bacterial density between control and exposed snails at any timepoint, or within cohorts over time, suggesting that bacterial density within the host hemolymph remains relatively stable. The snail host seems to maintain a relatively tight control over its bacteria, which schistosome infection does not significantly disturb.

Our results contrast with those obtained from bivalves infected with pathogens [9], where mollusk microbiomes are significantly disturbed. However, those studies involved transient pathogens that do not establish long-term relationships with their hosts and do not rely on host fitness for transmission. In cases of co-dependence, the impact on microbiomes can differ, though examples of this “strategy” in invertebrates are scarce. The impact of the parasitic snail *Coralliophila violacea* on the microbiome of its coral host, *Porites cylindrica,* offers an interesting parallel [41]. This sedentary snail employs a feeding strategy that minimizes tissue damage for its coral host, disturbing the coral microbiome only at the feeding site. In contrast, other mobile predatory snails tend to disturb coral microbiome on a larger scale. The limited effect of parasites on host microbiomes could actually be under strong natural selection, as parasite fitness is closely dependent on host fitness.

### 2. Stability of the microbiomes over time

The microbiomes of the snail hemolymph and hepatopancreas remained relatively stable over time. The weighted UniFrac distances showed no difference between hemolymph microbiomes across timepoints, and only a few inconsistencies in the hepatopancreas, with not specific patterns observed. Moreover, these differences became apparent only when using Bray-Curtis distances. This strongly suggests that the observed differences were likely due to small changes among closely related taxa, as the UniFrac distance accounts for the phylogeny of the microbiome community, unlike the Bray-Curtis distance. Additionally, microbiome diversity over time did not appear to be affected by the environmental variations: the high variability in microbial diversity in the water did not impact the diversity of the snail hemolymph or hepatopancreas microbiomes.

The apparent stability of host microbiomes may be attributed to survivorship bias, but it could also have a biological origin. Parasite infection negatively impacts the survival of snails (SuppFigure1), meaning that only the surviving snails are sampled. However, the survival of the infected snails during the pre-patent period was similar to that of the control snails, and decreased sharply only during the patent period. Therefore, any observation made during the pre-patent period should only be minimally impacted by this bias. Alternatively, the minimal disturbance in the host microbiome could result from parasite action or microbiome resilience. Schistosomes and their snail hosts have co-evolved for over 200 million years [18], and their fates are intimately linked during a relatively long developmental period of several weeks. Parasites have likely been under strong selective pressure to reduce their impact on host microbiomes to ensure their fitness and transmission. They may actively control their host microbiome using mechanisms such as extracellular vesicles [42,43] or may passively avoid disturbing the host microbiome by not triggering the immune system or by taking control of it. Interestingly, the immune status of infected snails during schistosome development, especially during the patent period, has received little attention. For example, phenoloxidase activity (a component of the humoral immune response in invertebrates) does not differ between infected and non-infected snails before week 7 (D49) post-infection, suggesting minimal impact of the parasite during the early stages of the patent period [21]. The minimal disturbance we observed could also be independent of the interaction between the host and parasite, instead explained by the resilience of the microbial community, which contributes to maintaining homeostasis within the host [44]. The snail tissue microbiomes are diverse [15], which can enhance the stability of the microbiome when challenged by stressful conditions [45]. Similarly, in oysters, which also exhibit diverse and distinct tissue microbiomes, changes in the hemolymph microbiome due to environmental stressors are limited to specific taxa [46].

### 3. What would be the consequences of manipulating the snail host microbiome for schistosome parasites?

If parasites do not disturb their host microbiomes to ensure their fitness and transmission, what would be the consequence of manipulating the host microbiome on the parasites? A limited number of studies has explored this topic. For instance, snails first treated with antibiotics and then exposed to schistosomes showed a shorter pre-patent period but produce fewer cercariae [47]. Antibiotic treatment of schistosome-infected snails led to the suppression of cercarial production [48]. While these results may suggest a complex interaction between the snail microbiome and the schistosome parasite, it is challenging to discern whether the impact on schistosome development was due to changes in the microbiome, or the effects of the antibiotics themselves. The use of axenic (germ-free) snails, generated without the administration of antibiotics [3], could help elucidate the role of the host microbiome in the interactions between snails and schistosomes. Absence of snail microbiome, which does not prevent snail infection [49], may affect schistosome fitness. The addition of a synthetic microbial community to axenic snails prior to or after schistosome infection may also reveal specific interactions.

### 4. Limitations

Our study has several limitations. First, our sample sizes per group, with 6-8 snails per cohort at each timepoint, were relatively small and may have limited our ability to detect subtle effects of parasitism on the host microbiomes. Second, we sampled different individuals from a snail population over time rather than resampling the same individuals. While we selected individuals from an inbred snail population [30,31], we cannot exclude the potential effect of snail genotypes on our measurements, which could have masked small effects of the parasites on the host microbiomes as well. Third, we used a single host-parasite combination in these experiments: further replication with different snail-schistosome combinations is needed to test the robustness of our findings and their possible generalization. For instance, Portet *et al.* did not find differences at day 25 (close to day 21 in our system) using SmBRE parasites and BgBRE snails [24]. While they used whole snail instead of hemolymph or hepatopancreas, SmBRE is also known to produce lower numbers of cercariae compared to SmLE parasites [31]. This could explain the absence of differences observed with that specific population of schistosome parasites. Fourth, a recent study has shown that miracidial stage may harbor a microbiome [50]. This parasite microbiome was not investigated in our study. However, it is unlikely that the presence of this parasite microbiome could significantly impact the microbiomes of the host, as we have shown no large impact of schistosome infection on the microbiome of the snail tissues in the pre-patent period. Finally, the snail environment was quite controlled: water and lettuce were changed daily, and lettuce was the only source of food. Clean water may also have limited the exchange of microbes between individuals. Different rearing conditions may reveal varying impacts of parasites on the microbiomes of their hosts.

## Supporting information

SuppFigure1

SuppFigure2

SuppFigure3

SuppFigure4

SuppFigure5

SuppTable1

SuppTable2

SuppTable3

## DECLARATIONS

### Ethics approval and consent to participate

This study was performed in accordance with the Guide for the Care and Use of Laboratory Animals of the National Institutes of Health. The protocol was approved by the Institutional Animal Care and Use Committee of Texas Biomedical Research Institute (permit number: 1419-MA).

### Consent for publication

Not applicable

### Availability of data and materials

Raw sequencing data are accessible from the NCBI Sequence Read Archive under BioProject accession number PRJNA1171869. Commands and scripts used for processing sequencing data and performing downstream analysis are available in a Jupyter notebook on Github (https://github.com/snordmeyer/S.mansoni-infected-B.glabrata-microbiomes).

### Authors’ contributions

SCN, TJCA, FDC, and WL designed the experiments. SCN prepared and collected the samples, extracted DNA and prepared the 16S rRNA gene libraries and performed the qPCR assays. SCN performed all the data analyses under FDC and WL supervision. SCN and FDC wrote the first version of the manuscript. All authors reviewed the manuscript and approved its final version.

## Acknowledgements

We thank Kathrin S. Jutzeler, Amanda Strickland, Robbie Diaz, Madison Morales and Neal Platt for snail and schistosome lifecycle maintenance.

## Supplementary material

**Supplementary Table 1. Average number of reads**

**Supplementary Table 2. Mean alpha-diversity over time**

**Supplementary Table 3. Alpha-Diversity statistics**

**Supplementary Figure 1. Survival Curve.**

**Supplementary Figure 2. Average Reads.**

**Supplementary Figure 3. Rarefaction Curves.**

**Supplementary Figure 4. Beta-Diversity: Homogeneity of Variances.**

**Supplementary Figure 5. Beta-Diversity: Additive models *day* comparisons.**

## References

1. Ooi MC, Goulden EF, Smith GG, Bridle AR. Haemolymph microbiome of the cultured spiny lobster Panulirus ornatus at different temperatures. Nature Scientific Reports [Internet]. 2019; Available from: 10.1038/s41598-019-39149-7

2. Stevens EJ, Bates KA, King KC. Host microbiota can facilitate pathogen infection. PLoS Pathog [Internet]. 2021;17. Available from: 10.1371/journal.ppat.1009514

3. Sun X, Hong J, Ding T, Wu Z, Lin D. Snail microbiota and snail–schistosome interactions: axenic and gnotobiotic technologies. Trends Parasitol [Internet]. 2024;40:241–56. Available from: 10.1016/j.pt.2024.01.002

4. Zheng D, Liwinski T, Elinav E. Interaction between microbiota and immunity in health and disease. Cell Res [Internet]. 2020;30:492–506. Available from: 10.1038/s41422-020-0332-7

5. Buffie CG, Pamer EG. Microbiota-mediated colonization resistance against intestinal pathogens. Nat Rev Immunol [Internet]. 2013;13:790–801. Available from: 10.1038/nri3535

6. Ubeda C, Djukovic A, Isaac S. Roles of the intestinal microbiota in pathogen protection. Clin Transl Immunology [Internet]. 2017;6:e128. Available from: 10.1038/cti.2017.2

7. Shi H, Yu X, Cheng G. Impact of the microbiome on mosquito-borne diseases. Protein Cell [Internet]. 2023;14:743–61. Available from: 10.1093/procel/pwad021

8. Beatriz A, Barletta F, Trisnadi N, Luis Ramirez J, Barillas-Mury C. Mosquito midgut prostaglandin release establishes systemic immune priming. iScience [Internet]. 2019;19:54–62. Available from: 10.1016/j.isci.2019.07.012

9. Destoumieux-Garzón D, Montagnani C, Dantan L, Nicolas NDS, Travers MA, Duperret L, et al. Cross-talk and mutual shaping between the immune system and the microbiota during an oyster’s life. Philosophical Transactions of the Royal Society B: Biological Sciences [Internet]. 2024;379. Available from: 10.1098/rstb.2023.0065

10. Leung JM, Graham AL, Knowles SCL. Parasite-microbiota interactions with the vertebrate gut: Synthesis through an ecological lens. Front Microbiol [Internet]. 2018;9. Available from: 10.3389/fmicb.2018.00843

11. Peachey LE, Jenkins TP, Cantacessi C. This gut ain’t big enough for both of us. Or is it? Helminth–microbiota interactions in veterinary species. Trends Parasitol [Internet]. 2017;33:619–32. Available from: 10.1016/j.pt.2017.04.004

12. Slatko BE, Luck AN, Dobson SL, Foster JM. Wolbachia endosymbionts and human disease control. Mol Biochem Parasitol [Internet]. 2014;195:88–95. Available from: 10.1016/j.molbiopara.2014.07.004

13. Telleria EL, Martins-da-Silva A, Tempone AJ, Traub-Cseko YM. Leishmania, microbiota and sand fly immunity. Parasitology [Internet]. 2018;145:1336–53. Available from: 10.1017/S0031182018001014

14. Chevalier FD, Diaz R, Mcdew-White M, Anderson TJ, Le Clec’h W. The hemolymph of Biomphalaria snail vectors of schistosomiasis supports a diverse microbiome. Environmental Microbiology [Internet]. 2020;22:5450–66. Available from: 10.1111/1462-2920.15303

15. Carruthers L V, Nordmeyer SC, Anderson TJ, Chevalier FD, Le Clec’h W. How should we sample snail microbiomes? 2024; Available from: 10.1101/2024.06.11.598555

16. Loker ES, Adema CM. Schistosomes, echinostomes and snails: comparative immunobiology. Parasitology Today [Internet]. 1995;11:120–4. Available from: 10.1016/0169-4758(95)80174-X

17. Pathak CR, Luitel H, Kjersti, Utaaker S, Khanal · Prabhat. One-health approach on the future application of snails: a focus on snail-transmitted parasitic diseases. Parasitol Res [Internet]. 2024;123:28. Available from: 10.1007/s00436-023-08021-z

18. Pila EA, Li H, Hambrook JR, Wu X, Hanington PC. Schistosomiasis from a snail’s perspective: advances in snail immunity. Trends Parasitol [Internet]. 2017;33:845–57. Available from: 10.1016/j.pt.2017.07.006

19. Verjee MA. Schistosomiasis: still a cause of significant morbidity and mortality. Res Rep Trop Med [Internet]. 2019; Available from: 10.2147/RRTM.S204345

20. Nelwan ML. Schistosomiasis: life cycle, diagnosis, and control. Current Therapeutic Research [Internet]. 2019;91:5–9. Available from: 10.1016/j.curtheres.2019.06.001

21. Le Clec’h W, Anderson TJC, Chevalier FD. Characterization of hemolymph phenoloxidase activity in two Biomphalaria snail species and impact of Schistosoma mansoni infection. Parasit Vectors [Internet]. 2016;9:32. Available from: 10.1186/s13071-016-1319-6

22. Meuleman EA, Holzmann PJ, Peet RC. The development of daughter sporocysts inside the mother sporocyst ofSchistosoma mansoni with special reference to the ultrastructure of the body wall. Zeitschrift für Parasitenkunde Parasitology Research [Internet]. 1980;61:201–12. Available from: 10.1007/BF00925512

23. Castillo MG, Humphries JE, Mourão MM, Marquez J, Gonzalez A, Montelongo CE. Biomphalaria glabrata immunity: Post-genome advances. Dev Comp Immunol [Internet]. 2020;104. Available from: 10.1016/j.dci.2019.103557

24. Portet A, Toulza E, Lokmer A, Huot C, Duval D, Galinier R, et al. Experimental infection of the biomphalaria glabrata vector snail by schistosoma mansoni parasites drives snail microbiota dysbiosis. Microorganisms [Internet]. 2021;9. Available from: 10.3390/microorganisms9051084

25. Silva TM, Melo ES, Lopes ACS, Veras DL, Duarte CR, Alves LC, et al. Characterization of the bacterial microbiota of biomphalaria glabrata (Say, 1818) (Mollusca: Gastropoda) from Brazil. Lett Appl Microbiol [Internet]. 2013;57:19–25. Available from: 10.1111/lam.12068

26. Allan ERO, Tennessen JA, Sharpton TJ, Blouin MS. Allelic variation in a single genomic region alters the microbiome of the snail Biomphalaria glabrata. Journal of Heredity [Internet]. 2018;604–9. Available from: 10.1093/jhered/esy014

27. Huot C, Gerardo NM, Spor A, Novakova E, Clerissi C, Gourbal B, et al. Schistosomiasis vector snails and their microbiota display a phylosymbiosis pattern. Front Microbiol [Internet]. 2020;10. Available from: 10.3389/fmicb.2019.03092

28. Schols R, Vano Verber Ghe I, Huyse T, Decaestecker E. Host-bacteriome transplants of the schistosome snail host Biomphalaria glabrata reflect species-specific associations. FEMS Microbiol Ecol [Internet]. 2023;99:1. Available from: 10.1093/femsec/fiad101

29. McCann P, McFarland C, Megaw J, Siu-Ting K, Cantacessi C, Rinaldi G, et al. Assessing the microbiota of the snail intermediate host of trematodes, Galba truncatula. Parasit Vectors [Internet]. 2024;17. Available from: 10.1186/s13071-024-06118-7

30. Bonner KM, Bayne CJ, Larson MK, Blouin MS. Effects of Cu/Zn superoxide dismutase (sod1) genotype and genetic background on growth, reproduction and defense in Biomphalaria glabrata. PLoS Negl Trop Dis [Internet]. 2012;6. Available from: 10.1371/journal.pntd.0001701

31. Le Clec’h W, Diaz R, Chevalier FD, McDew-White M, Anderson TJC. Striking differences in virulence, transmission and sporocyst growth dynamics between two schistosome populations. Parasit Vectors [Internet]. 2019;12. Available from: 10.1186/s13071-019-3741-z

32. Hornung BVH, Zwittink RD, Kuijper EJ. Issues and current standards of controls in microbiome research. FEMS Microbiol Ecol [Internet]. 2019;95:45. Available from: 10.1093/femsec/fiz045

33. Apprill A, Mcnally S, Parsons R, Weber L. Minor revision to V4 region SSU rRNA 806R gene primer greatly increases detection of SAR11 bacterioplankton. Aquatic Microbial Ecology [Internet]. 2015;75:129–37. Available from: 10.3354/ame01753

34. Bolyen E, Rideout JR, Dillon MR, Bokulich NA, Abnet CC, Al-Ghalith GA, et al. Reproducible, interactive, scalable and extensible microbiome data science using QIIME 2. Nat Biotechnol [Internet]. 2019;37:852–7. Available from: 10.1038/s41587-019-0209-9

35. R Core Team. R: A language and environment for statistical computing. [Internet]. Vienna, Austria: R Foundation for Statistical Computing; 2021. Available from: https://www.R-project.org/

36. Beideman D, Frled B, Sherma J. Effects of Schistosoma mansoni infection on the survival, fecundity, and triacylglycerol content of Biomphalaria glabrata snails. J Vet Sci Mad Oiagn [Internet]. 2013;2:3. Available from: 10.4172/2325-9590.1000115

37. Crews AE, Yoshino TP. Schistosoma mansoni: Effect of infection on reproduction and gonadal growth in Biomphalaria glabrata. Exp Parasitol [Internet]. 1989;68:326–34. Available from: 10.1016/0014-4894(89)90114-8

38. Cooper LA, Larson SE, Lewis FA. Male reproductive success of Schistosoma mansoni-infected Biomphalaria glabrata snails. J Parasitol [Internet]. 1996;82(3):428–31. Available from: https://pubmed.ncbi.nlm.nih.gov/8636847/

39. Cortés A, Clare S, Costain A, Almeida A, McCarthy C, Harcourt K, et al. Baseline gut microbiota composition is associated with Schistosoma mansoni infection burden in rodent models. Front Immunol [Internet]. 2020;11. Available from: 10.3389/fimmu.2020.593838

40. Hu Y, Chen J, Xu Y, Zhou H, Huang P, Ma Y, et al. Alterations of gut microbiome and metabolite profiling in mice infected by Schistosoma japonicum. Front Immunol [Internet]. 2020;11. Available from: 10.3389/fimmu.2020.569727

41. Clements CS, Burns AS, Stewart FJ, Hay ME. Parasite-host ecology: the limited impacts of an intimate enemy on host microbiomes. Anim Microbiome [Internet]. 2020;2:42. Available from: 10.1186/s42523-020-00061-5

42. Rooney J, Northcote HM, Williams TL, Cortés A, Cantacessi C, Morphew RM. Parasitic helminths and the host microbiome – a missing ‘extracellular vesicle-sized’ link? Trends Parasitol [Internet]. 2022;38:737–47. Available from: 10.1016/j.pt.2022.06.003

43. Avni D, Avni O. Extracellular vesicles: schistosomal long-range precise weapon to manipulate the immune response. Front Cell Infect Microbiol [Internet]. 2021;11. Available from: 10.3389/fcimb.2021.649480

44. Philippot L, Griffiths BS, Langenheder S. Microbial community resilience across ecosystems and multiple disturbances. Microbiology and Molecular Biology Reviews [Internet]. 2021;85. Available from: 10.1128/mmbr.00026-20

45. Dogra SK, Doré J, Damak S. Gut microbiota resilience: definition, link to health and strategies for intervention. Front Microbiol [Internet]. 2020; Available from: 10.3389/fmicb.2020.572921

46. Dupont S, Lokmer A, Corre E, Auguet J, Petton B, Toulza E, et al. Oyster hemolymph is a complex and dynamic ecosystem hosting bacteria, protists and viruses. Anim Microbiome [Internet]. 2020;2:12. Available from: 10.1186/s42523-020-00032-w

47. Melchiorre HG, Gutierrez SO, Minchella DJ, Vannatta JT. Downstream effects: Impact of antibiotic pollution on an aquatic host–parasite interaction. Ecosphere [Internet]. 2023;14. Available from: 10.1002/ecs2.4513

48. Warren KS, Peters PA. Quantitative aspects of exposure time and cercarial dispersion on penetration and maturation of Schistosoma mansoni in mice. American Journal of Tropical Medicine and Hygiene [Internet]. 1967;16:718–22. Available from: 10.1080/00034983.1967.11686489

49. Chernin E. Infection of Australorbis glabratus with Schistosoma mansoni under bacteriologically sterile conditions. Exp Biol Med [Internet]. 1960;105:292–6. Available from: 10.3181/00379727-105-26088

50. Formenti F, Cortés A, Deiana M, Salter S, Parkhill J, Berriman M, et al. The human blood fluke, Schistosoma mansoni, harbors bacteria throughout the parasite’s life cycle. J Infect Dis [Internet]. 2023;228:1299–303. Available from: 10.1093/infdis/jiad288

