## Supplementary figures and images for "Limited Impact of Schistosome Infection on *Biomphalaria glabrata* Snail Microbiomes"

### SuppFigure1

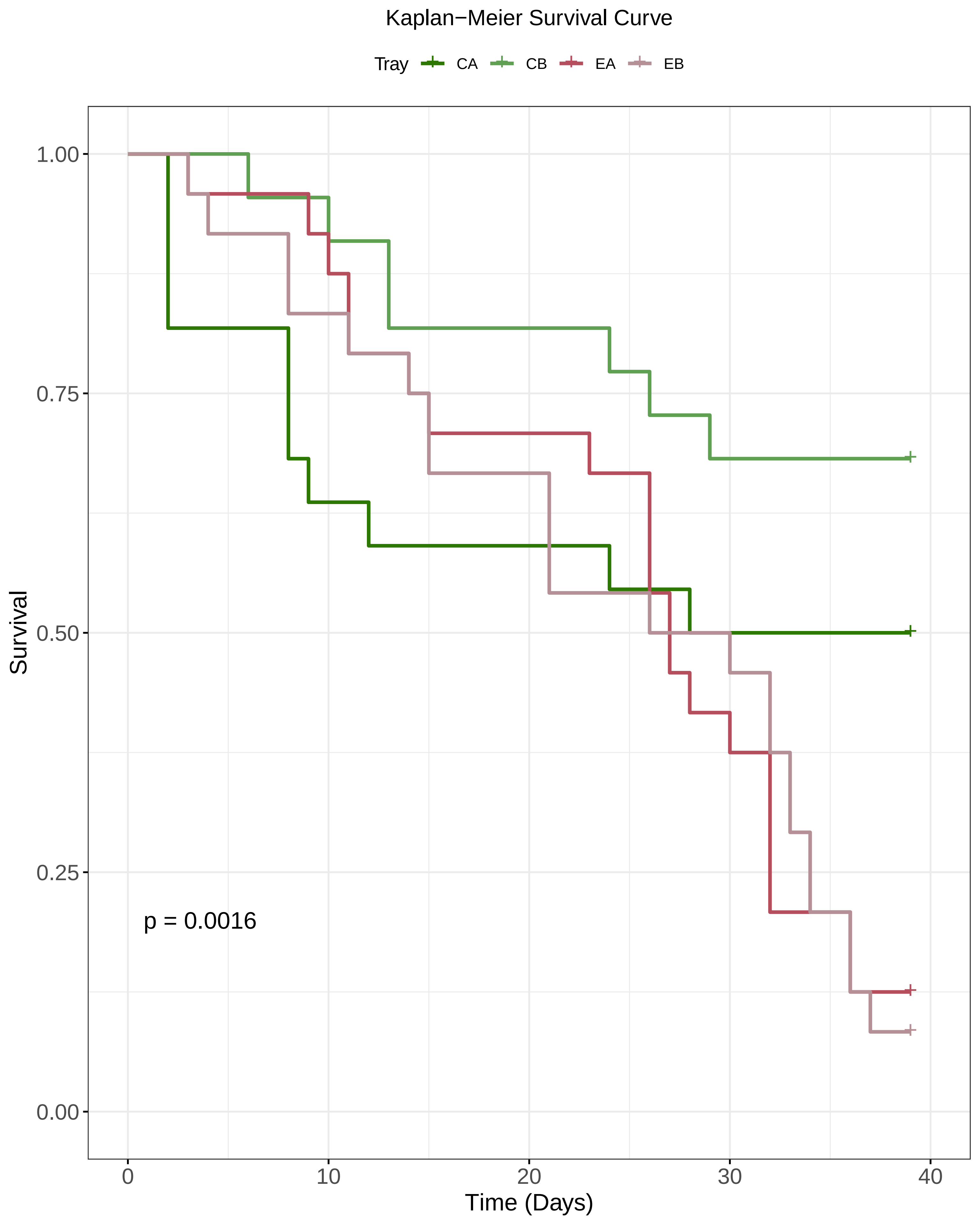

### SuppFigure2

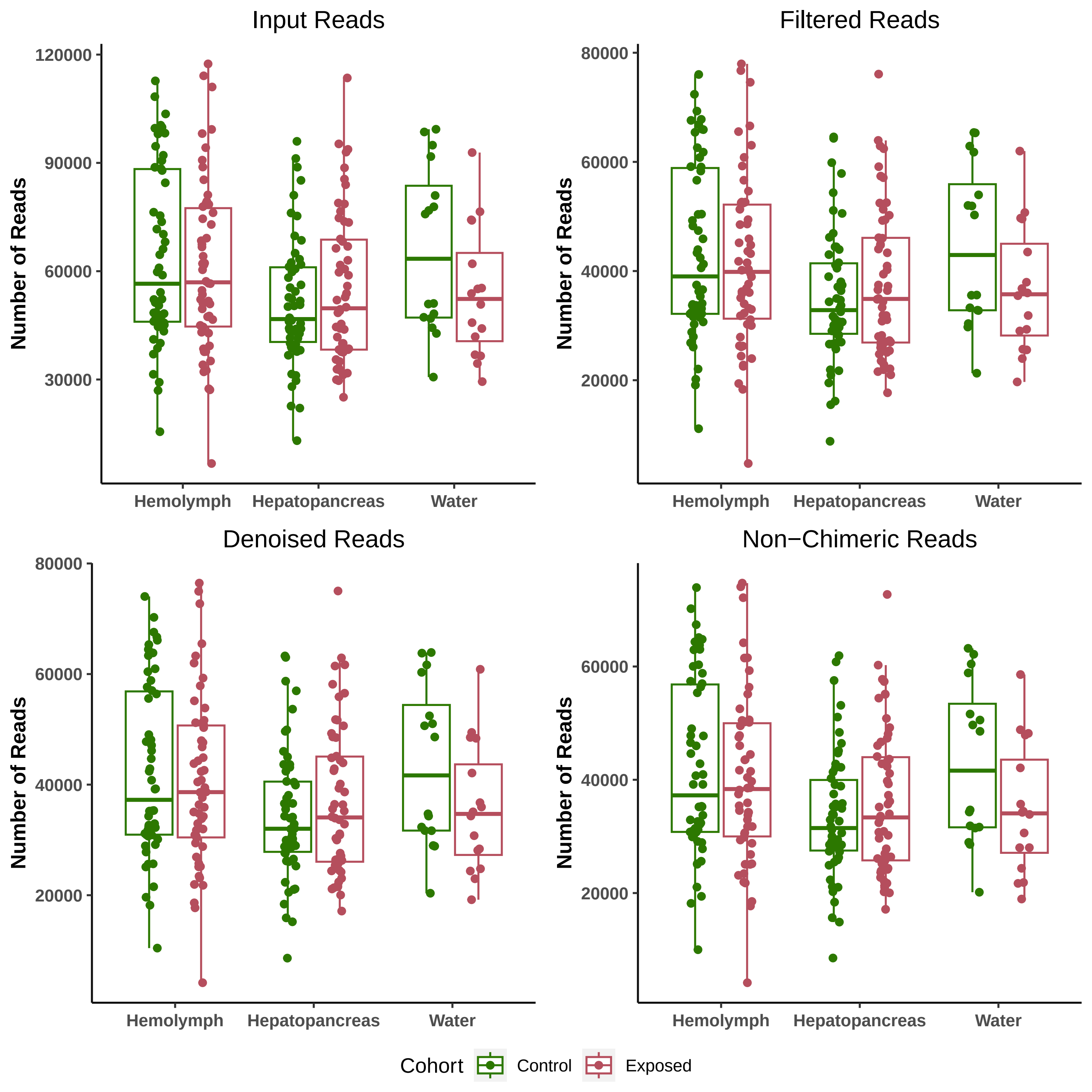

### SuppFigure3

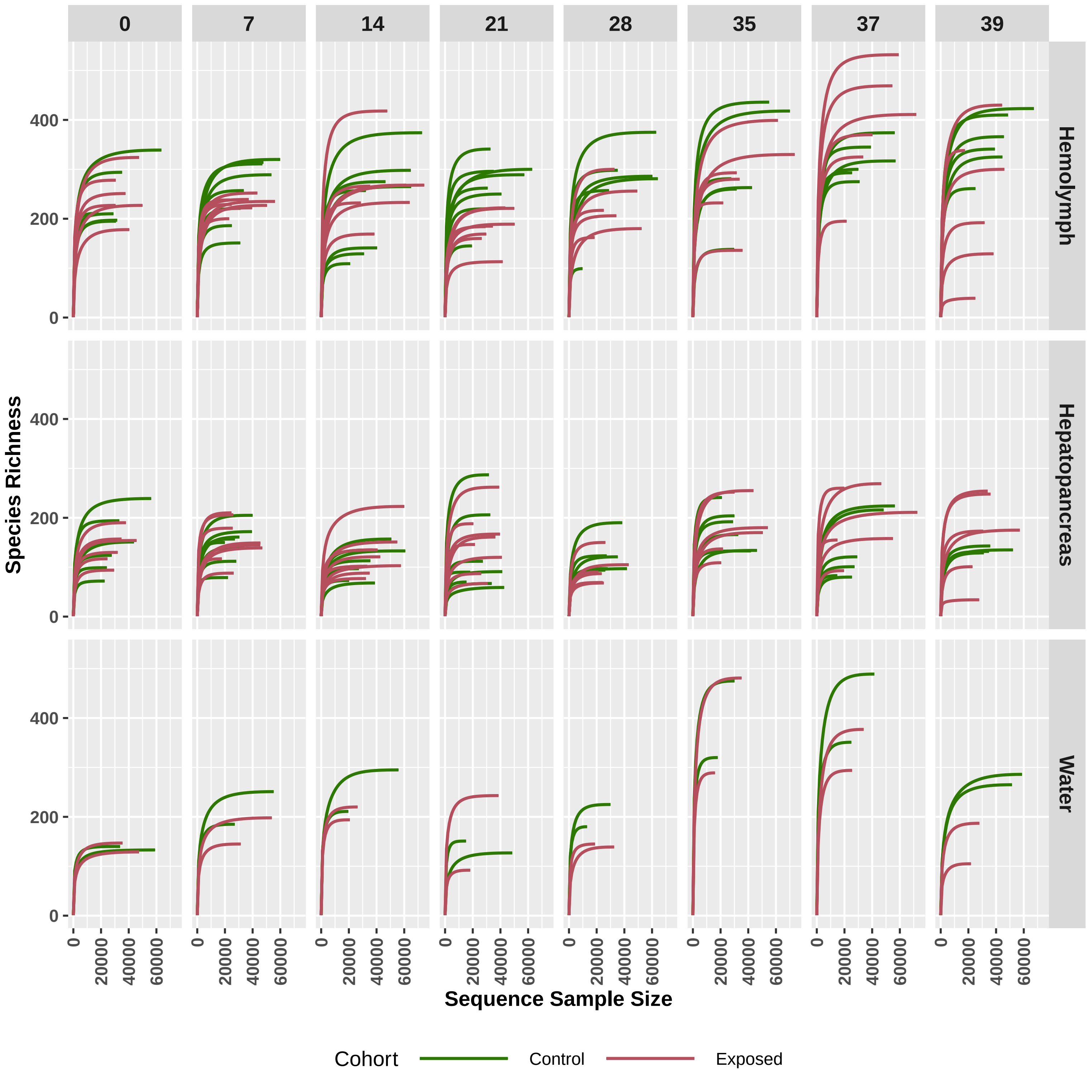

### SuppFigure4

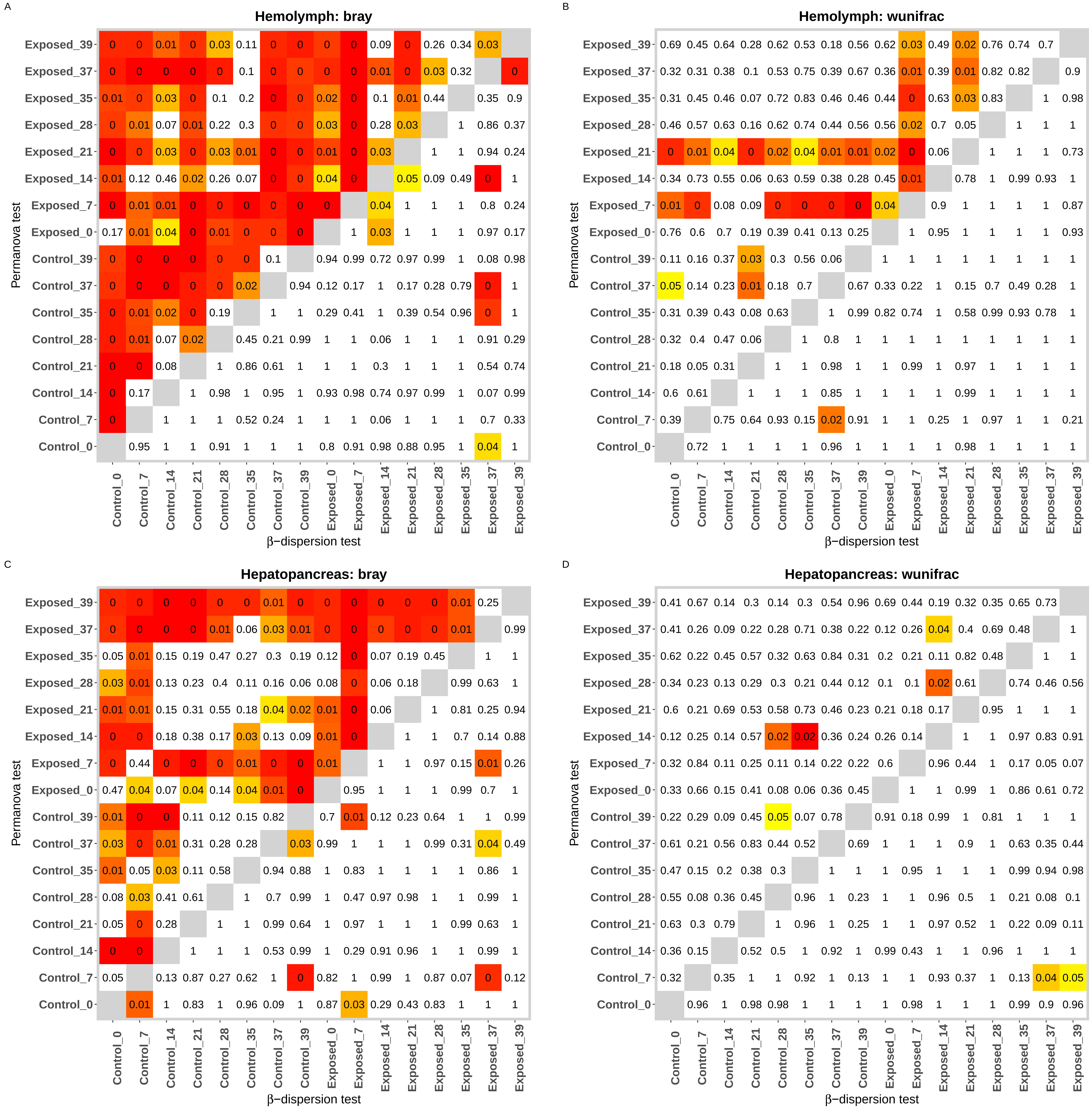

### SuppFigure5

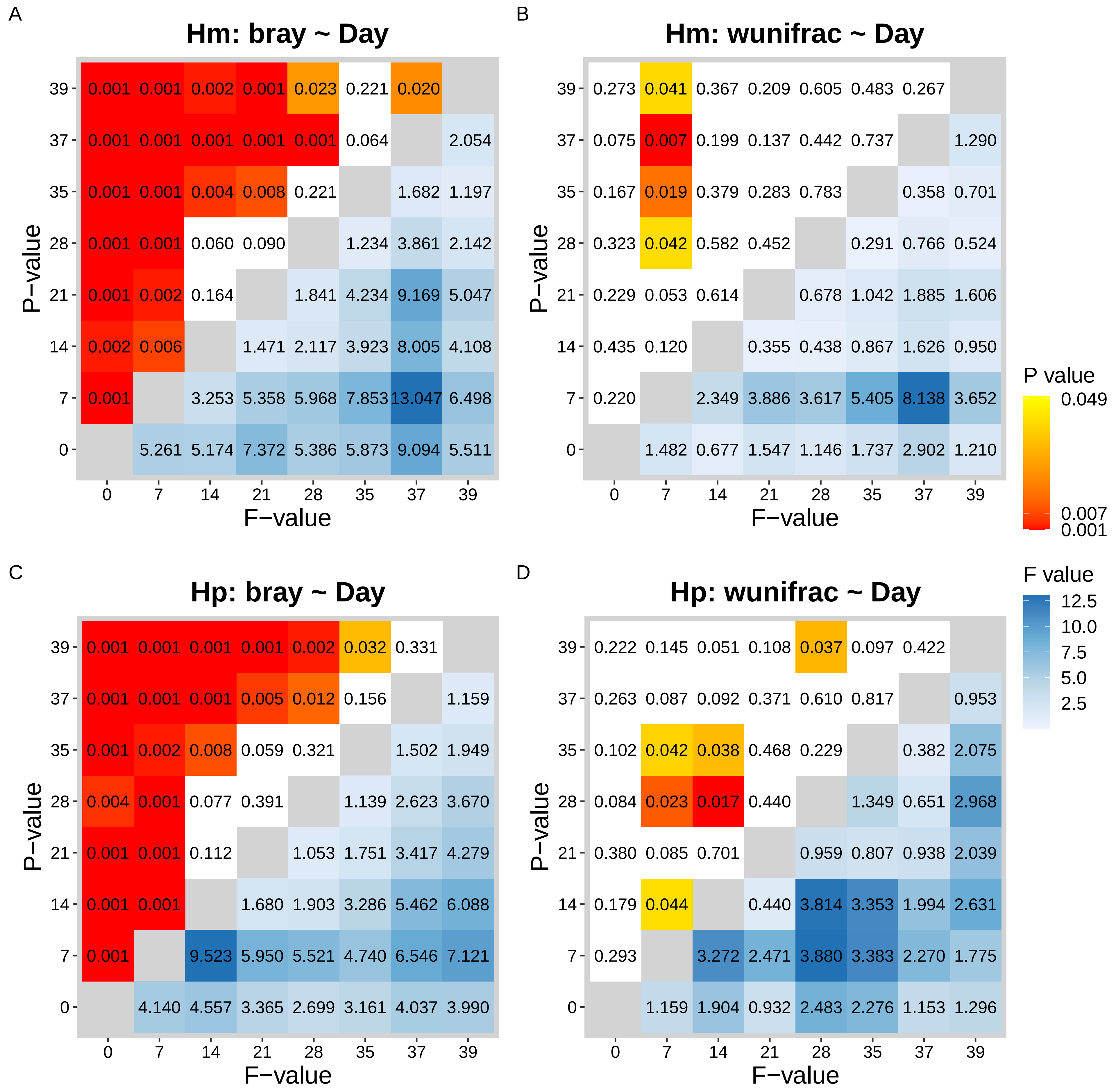
